# Habitat predicts levels of genetic admixture in *Saccharomyces cerevisiae*

**DOI:** 10.1101/095265

**Authors:** Viranga Tilakaratna, Douda Bensasson

## Abstract

Genetic admixture can provide material for populations to adapt to local environments, and this process has played a crucial role in the domestication of plants and animals. The model yeast, *Saccharomyces cerevisiae*, has been domesticated multiple times for the production of wine, sake, beer and bread, but the high rate of admixture between yeast lineages has so far been treated as a complication for population genomic analysis. Here we make use of the low recombination rate at centromeres to investigate admixture in yeast using a classic Bayesian approach and a more conservative locus by locus phylogenetic approach developed here. Using both approaches, we find that *S. cerevisiae* from stable oak woodland habitats are less likely to show recent genetic admixture compared with those isolated from transient habitats such as fruits, wine or human infections. When woodland yeast strains do show recent genetic admixture, the degree of admixture is lower than in strains from other habitats. Furthermore, *S. cerevisiae* populations from oak woodlands are genetically isolated from each other, with only occasional migration between woodlands and local fruit habitats. Application of our phylogenetic approach suggests that there is a previously undetected population in North Africa that is the closest outgroup to the European *S. cerevisiae*, including the domesticated Wine population. Thorough testing for admixture in *S. cerevisiae* therefore leads to a better understanding of the underlying population structure of the species and will be important for understanding the selective processes underlying domestication in this economically important species.

## Introduction

The wine yeast *Saccharomyces cerevisiae* is one of the most economically important model organisms and is used by humans around the world to produce alcohol and to ferment foods (Fay and Benavides, 2005; Wang *et al.*, 2012; Gallone *et al.*, 2016; Gonçalves *et al.*, 2016). *S. cerevisiae* is also found in the wild on fruits, flowers and on the bark of trees in oak woodlands and can occur as a commensal or pathogen of humans (Sniegowski *et al.*, 2002; Wang *et al.*, 2012; Hyma and Fay, 2013; Goddard *et al.*, 2010; Cromie *et al.*, 2013; Dashko *et al.*, 2016; Goddard and Greig, 2015). *S. cerevisiae* shares the oak woodland habitat with all other *Saccharomyces* species, suggesting that the woodland habitat is ancestral for the species (Eberlein *et al.*, 2015). The presence of *S. cerevisiae* on a broad range of habitats compared to its closest relatives makes it an ideal model for molecular ecology, especially because many genome-wide technologies are developed and tested first on *S. cerevisiae*, providing a wealth of supporting resources to help interpret ecological patterns (Cherry *et al.*, 2011).

However, the presence of genetic admixture among natural strains of yeast presents a challenge for the use of *S. cerevisiae* as a model in population genomics (Liti *et al.*, 2009; Almeida *et al.*, 2015; Ludlow *et al.*, 2016; Barbosa *et al.*, 2016). Indeed, genetic admixture also complicates the population genomic analysis of model plants (Hufford *et al.*, 2013; Brandvain *et al.*, 2014) and animals (Pool *et al.*, 2012), including humans (Sankararaman *et al.*, 2014; Harris and Nielsen, 2016). In addition to informing population genomic analysis, the study of genetic admixture can also reveal signatures of selective processes in natural (Brandvain *et al.*, 2014) and human commensal populations (Hufford *et al.*, 2013; Pool *et al.*, 2012). Introgressions from natural to domesticated populations can allow adaptation of crops to local habitats (Hufford *et al.*, 2013) and has probably played an important role in animal domestication (Marshall *et al.*, 2014). The study of genetic admixture with deleterious effects in natural populations also has potential applications in conservation biology (Harris and Nielsen, 2016). In the case of *S. cerevisiae*, analysis of genetic admixture could potentially reveal mechanisms of adaptation to industrial applications and the human body, as well as the connectivity of natural populations within and between habitats.

Past studies have employed a number of different approaches to test whether strains are “mosaics” (genetically admixed), which precludes comparison among studies or samples (Liti *et al.*, 2009; Almeida *et al.*, 2015; Wang *et al.*, 2012; Cromie *et al.*, 2013; Barbosa *et al.*, 2016). The precise definitions differ among studies, but in general, admixed *S. cerevisiae* strains are identified as (i) those that have long branches in phylogenetic analyses and do not occur in well-supported clades with other strains (Liti *et al.*, 2009; Wang *et al.*, 2012), or (ii) those that are not assigned to distinct populations using a Bayesian clustering method (Liti *et al.*, 2009; Almeida *et al.*, 2015; Wang *et al.*, 2012; Cromie *et al.*, 2013; Barbosa *et al.*, 2016). In all cases, conclusions are based on analysis of pooled genome data or pooled data from multiple loci. The major drawback of using pooled data to test for similarity to well-sampled populations is that strains belonging to populations that are poorly represented in a sample may incorrectly be defined as mosaics.

Here we test for differences in the levels of admixture among *S. cerevisiae* found in different habitats using two different approaches. We employ the most commonly used Bayesian test to detect admixture in our study strains (Pritchard *et al.*, 2000), and we also develop a more conservative locus by locus phylogenetic approach to test for admixture on every chromosome. We avoid the complicating effects of selection and recombination in our analysis by using sequences for the point centromeres of *S. cerevisiae*, which are easy to sequence, neutrally and rapidly evolving and have low recombination rates (Bensasson *et al.*, 2008). Using complete DNA sequence for the centromeres and the flanking DNA of all 16 chromosomes from 80 *S. cerevisiae* strains, we show differences between habitats in levels of genetic admixture and that oak woodland populations are more isolated than those from other habitats.

## Materials and Methods

### Yeast strains and DNA sequencing

We analyzed DNA sequence data from all 16 centromeres for 80 *S. cerevisiae* strains (Table 1 and Supplemental File 1). Centromere sequence was already available for 33 strains (Table 1; Bensasson, 2011; GenBank: HQ339369-HQ339877), and we re-use these data here. These previously-reported sequences were obtained from monosporic derivatives, which we expect to be completely homozygous in all parts of the genome except at the MAT locus (Bensasson, 2011).

**Table 1:**
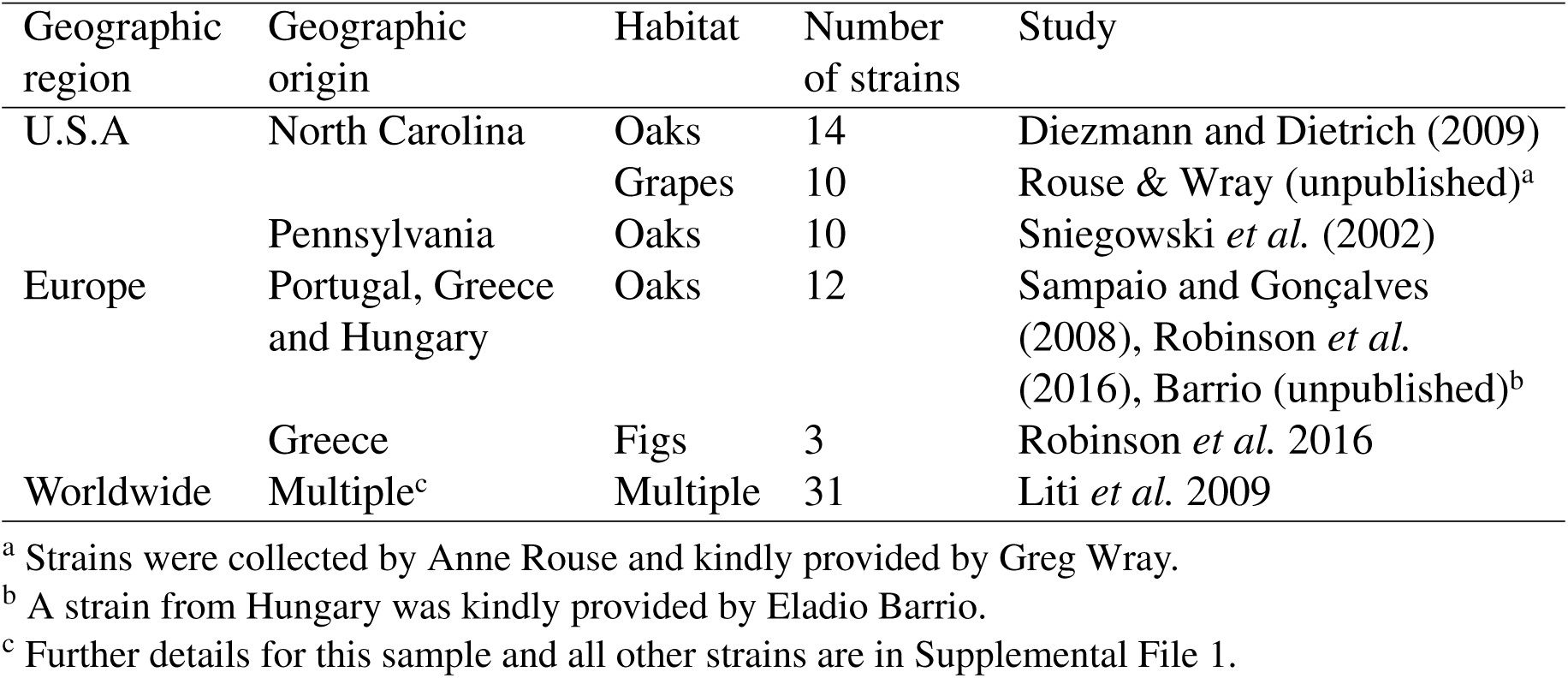
Summary of the 80 *S. cerevisiae* strains used in this study.

For the remaining 47 strains, we generated monosporic derivatives by sporulating yeast and isolating single spores as described in Amberg *et al.* (2005). DNA was extracted, amplified and sequenced from the monosporic derivatives using the extraction, PCR and DNA sequencing conditions described in Bensasson (2011). DNA sequence reads were assembled into a consensus DNA sequence for each strain at each locus using Staden version 1.7.0 (Bonfield *et al.*, 1995) and its quality was assessed using Phred (version: 0.020425.c) as described in Bensasson (2011). Low quality bases at the ends of consensus sequences were trimmed and any other bases with a Phred-scaled quality score below q40 were masked. The methods used here and the q40 filter ensure a very low base-calling error rate (Bensasson, 2011). The resulting centromere sequences are available in GenBank (KT206234-KT206982). Sequences were manually aligned and visualised in SeaView 4.0 (Gouy *et al.*, 2010).

### Phylogenetic analysis

Alignments for all 16 centromere loci were concatenated into a single long alignment using a custom perl script (alcat.pl). Genetic distances between DNA sequences were estimated and analyzed using the ape package (version 3.5 Paradis, 2011) in R (version 3.3.0). More specifically, we estimated genetic distance using the F84 model (Felsenstein and Churchill, 1996) as implemented in dist.dna and constructed a neighbour joining tree (Saitou and Nei, 1987) from these distances using nj (Paradis, 2011). DNA sequence data were bootstrapped using boot.phylo with 10,000 replicates to test the statistical support of the clades obtained. The resulting phylogram with associated bootstrap values was visualised and coloured using plot.phylo (Figure 2).

**Figure 1:**
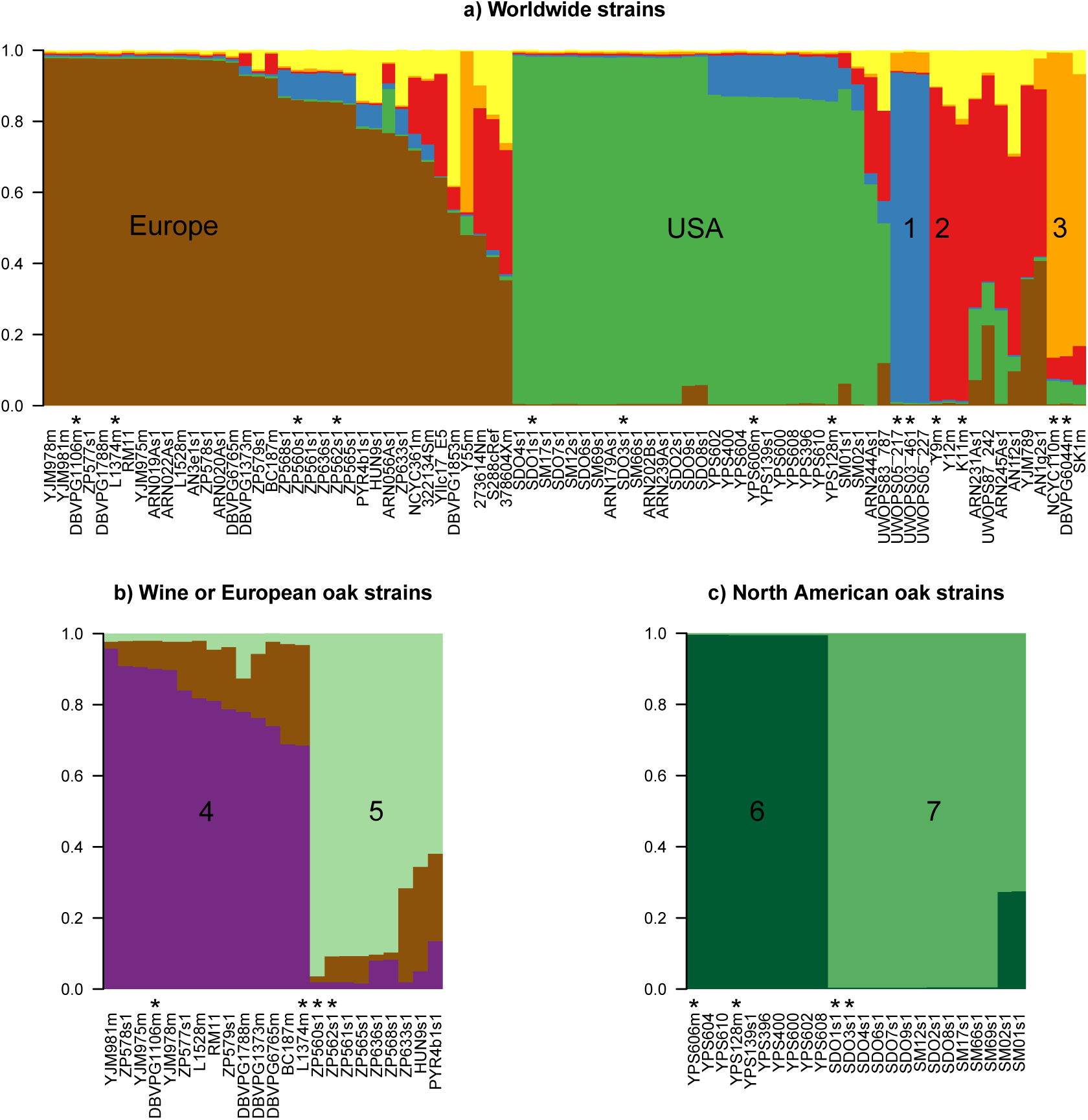
Identification of seven *S. cerevisiae* populations when including the subpopulations within Europe and the USA. A breakdown of population assignments defined by the most likely models estimated by *structure* analysis. Reference strains for the detection of admixture in subsequent analyses are highlighted with a “*”. **a) Five major global populations identified in the worldwide sample.** Most of our worldwide sample of 80 strains can be assigned to 5 distinct populations: 1 from Malaysia (blue), 2 from sake (red), and 3 from West Africa (orange), as well as strains from Europe (brown) and the USA (green). *structure* also invoked some gene flow from a sixth (K=6) population (yellow), however we did not encounter any strains that clearly represent this population. **b) Two subpopulations identified in Europe.** Analysis of the 23 strains that were isolated from European oak trees or were previously assigned to a European Wine population (Liti *et al.*, 2009) shows that there are at least two *S. cerevisiae* subpopulations in Europe (K=3): population 4 from wine (purple), 5 from European Oaks (green). **c) Two subpopulations identified in the USA.** Analysis of the 24 strains isolated from oaks in the USA shows that there are two *S. cerevisiae* subpopulations in the USA (K=2): population 6 from Pennsylvanian Oaks (dark green), and population 7 from North Carolina Oaks (light green).

**Figure 2:**
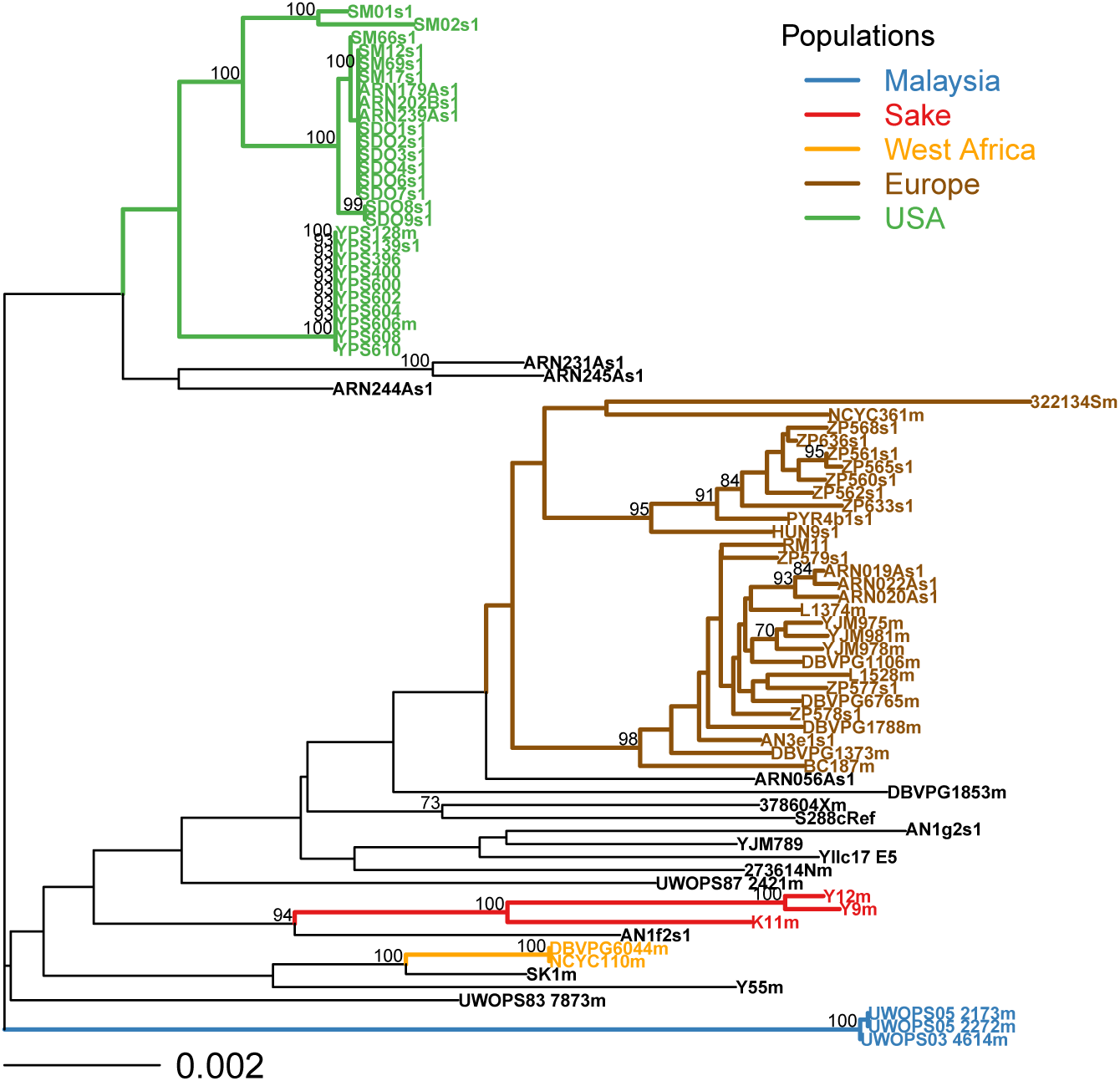
Neighbour joining distance analysis of all strains shows subpopulations within the USA and Europe. Bootstrap support was estimated from 10,000 bootstrap replicates and support is shown as a percentage for clades with over 70% support. Strain names and clades are coloured according to populations defined by the *structure* analysis shown in Figure 1a. Bootstrap support for the European and USA populations is lower than 70%, however there is strong bootstrap support (at least 95%) for subpopulations within the USA and Europe.

A second phylogenetic analysis was performed, after excluding strains showing recent genetic admixture (Supplemental File 1). We conducted a phylogenetic analysis of the concatenated alignment of data for all loci using the maximum likelihood approach implemented in RAxML (version 8.2.4; Stamatakis, 2014). We used a general time reversible model with a gamma distribution to estimate rate heteregeneity at sites from the data (GTRGAMMA), and a rapid bootstrap analysis for 10,000 bootstrap replicates with a search for the best-scoring Maximum Likelihood tree in the same RAxML run (Figure 3).

**Figure 3:**
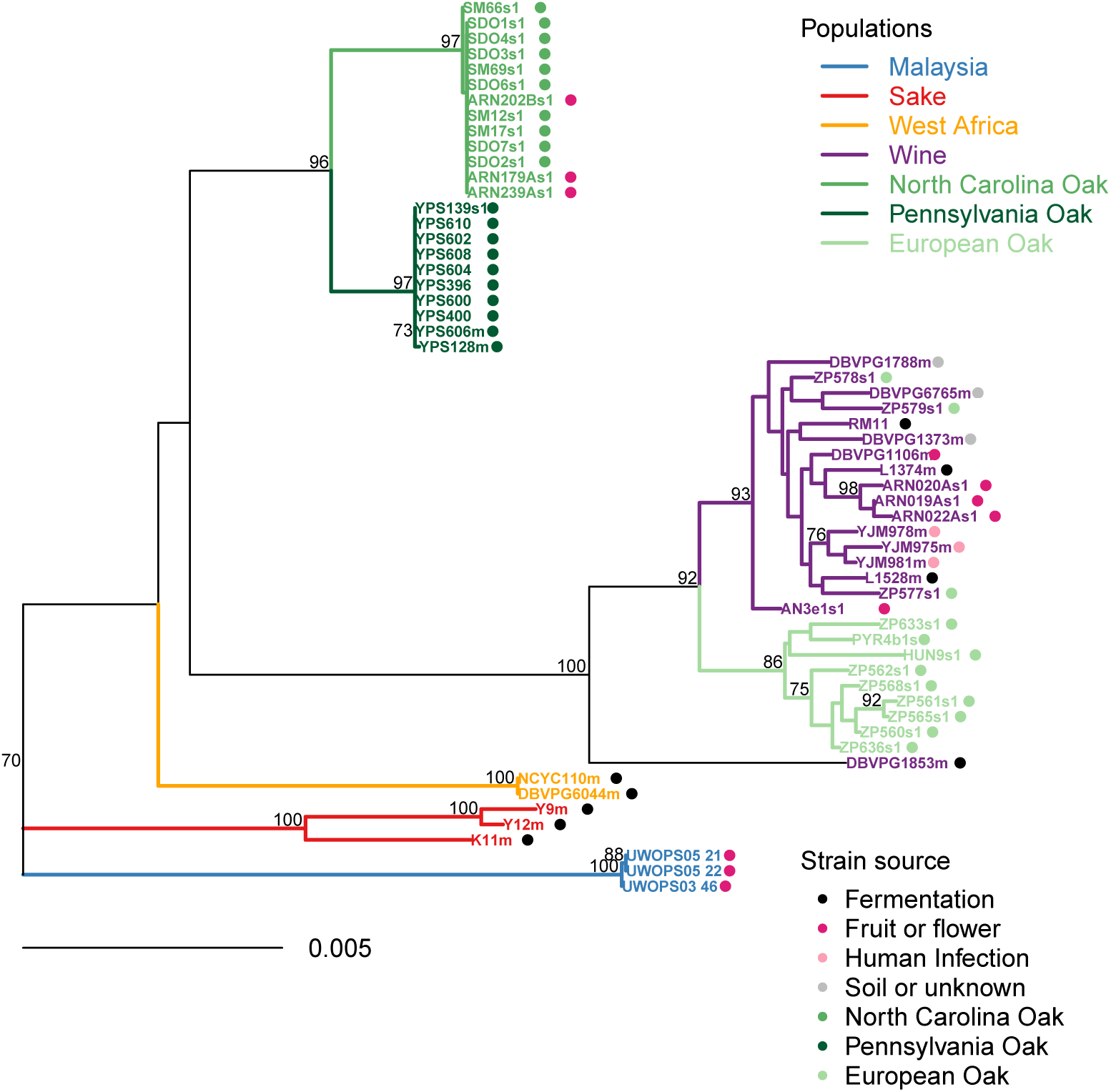
Phylogenetic analysis of non-admixed strains reveals structured populations with occasional migration. We excluded 22 recently admixed strains out of the 80 strains in the original dataset, and estimated the phylogeny using a maximum likelihood approach with bootstrap support shown for clades with over 70% support out of 10,000 bootstrap replicates. Clades are coloured according to their genetic population, and strains are annotated according to their habitat. DBVPG1853m, which is not clearly assigned to a European population in this maximum likelihood analysis appeared most similar to the Wine population in the locus by locus analysis. Strains isolated from oak trees are similar to strains isolated from the same woodland and distinct from those isolated from other regions. There is some migration of *S. cerevisiae* from the North Carolina Oak population onto North Carolina grapes (prefixed “ARN”). In addition, all three strains isolated from oak trees in Aldeia das Dez in Portugal (prefixed “ZP57”) had migrated from the Wine population to trees.

### Population structure analysis

We tested for population structure within our sample of 80 yeast strains using the software package *structure* (version 2.3.4) with a model taking account of linkage between polymorphisms at the same locus (Falush *et al.*, 2003), and assuming the DNA sequences were haploid. Using *structure*, we estimated the most likely number of populations to explain the data (K) by varying K from 1 to 10 and visualizing the results. The linkage model allows individuals to show admixture between the K different populations (INFERALPHA 1), and we ran it with default parameters: a burnin of 10,000 steps followed by a run length of 20,000 steps. In pilot experiments we found that increasing the length of the burnin from 10,000 to 100,000 did not alter our conclusions. Runs were repeated for each value of K five times, and we identified the model with the highest log likelihood and lowest value of K across the 50 runs. In addition, we visually verified that larger values of K resulted in qualitatively similar results to those for the minimum values of K discussed in the Results.

The *structure* package was run using three perl scripts available at https://github.com/bensassonlab/scripts (i) structureInfile.pl converts sequence alignments (in fasta format) to *structure* input files that summarise bases at variable sites; (ii) structureShell.pl runs *structure* one time for each value of K in a specified range (from 1 to 10 in this study); (iii) structurePrint.pl plots the structure results as barplots using R and allows for user specification of colors.

Our preliminary phylogenetic analysis showed well-supported subpopulations within Europe and the USA (at least 95% bootstrap support for all 4 subpopulations in Figure 2) that are missed by the initial *structure* analysis (Figure 1a). Therefore we repeated the above *structure* analysis of 50 runs (K=1 to 10, 5 replicates) on two subsets of the data: (i) on the 23 strains that were either assigned to the European Wine population by Liti *et al.* (2009) or were isolated from European oak trees (Supplemental File 1, Figure 1b); and (ii) on the 24 strains isolated from oaks in the USA (Table 1, Figure 1c).

### Testing for recent genetic admixture

In order to test for genetic admixture between populations, we first defined distinct *S. cerevisiae* populations on the basis of *structure* and phylogenetic analyses (Figures 1 and 2). From each of the seven different populations identified in this way, we chose two reference strains to define each population (the minimum number that *structure* needs to estimate allele frequencies within a population). Where possible, reference strains were chosen from representative strains that were previously assigned to a particular population (Liti *et al.*, 2009). For example, for the Wine population, we chose strains DBVPG1106m and L1374m, which were previously assigned to the Wine population by Liti *et al.* (2009) and were isolated from wine. For the remaining populations, European Oak and North Carolina Oak, reference strains were those with the highest estimates of genetic ancestry for their respective population (Figure 1b and 1c).

The remaining 66 strains in this study were assigned to populations or defined as showing recent genetic admixture using two independent approaches. The first approach uses *structure* to estimate levels of admixture for every strain with reference strains to define distinct populations using the USEPOPINFO option (Pritchard *et al.*, 2000). Preliminary analysis suggested that when *structure* was used in this manner it incorrectly invoked admixture to explain the genetic composition of strains that did not belong to the defined populations yet showed no evidence of admixture. Therefore, we developed a second locus by locus phylogenetic test for admixture that only classifies strains as admixed if they show well supported similarity to multiple populations, and is thus more conservative. Details of both approaches are described in the following sections.

#### (i) Detecting admixture using *structure*

We ran *structure* using the same parameters as we used for defining populations above, except that we used the USEPOPINFO model in *structure* (Pritchard *et al.*, 2000) to estimate ancestry for the 66 *S. cerevisiae* strains of unknown origin (POPFLAG=0) by pre-specifying the population of origin for the 14 reference strains (POPFLAG=1). We set the number of populations to seven (K=7), and selected the breakdown of ancestry components for each strain from the most likely model out of 20 independent runs. For consistency across our two approaches, we defined a strain as “admixed” if its proportion of ancestry to a single population was less than 0.94 (equivalent to 1 out of 16 centromere loci being from a different population).

#### (ii) Detecting admixture using a locus by locus phylogenetic approach

We also performed locus by locus phylogenetic analyses of all 80 strains, assigning each of the 66 non-reference strains at each locus to a population according to which of the 14 reference strains it grouped with. We used custom perl scripts to run the phylogenetic analysis using the ape package in R. For each locus, we constructed a neighbour joining tree of genetic distances using the F84 model, used 10,000 bootstrap replicates to assess statistical support for each clade and output a text summary of the strains found in each clade using the prop.part tool of boot.phylo in ape (Paradis, 2011). We only considered clades that had at least 70% bootstrap support. For each locus, strains found in clades with reference strains from only one population were assigned to the same population as the reference strains. Because of the limited phylogenetic resolution at some loci, we also assigned strains in clades with reference strains only from the European Oak or Wine populations as “European” and those in clades with North Carolina or Pennsylvania reference strains as “USA”. In cases where a sequence does not group with those of reference strains belonging to a single population, its population status at that locus is classed as “undefined”. The more general classifications of “European” or “USA” do not conflict with subpopulation classifications within those groups, and loci classed as “undefined” do not conflict with classifications at any other loci. We then compared population predictions across all loci for a strain, and if a strain was assigned to a population at a locus that conflicted with the population assignment at any of the other loci, then that strain was defined as showing genetic admixture. For example, SDO8s1 was isolated from a North Carolina oak, and has a CEN4 sequence from the Wine clade, but it has 5 loci that are in the same clade as the North Carolina Oak reference strains (the remaining loci were undefined or in the USA clade; Figure S1). This strain therefore shows admixture between Wine and North Carolina Oak populations.

In most cases (55 out of 66 strains) the two approaches resulted in the same population and admixture assignments. Fewer strains were defined as admixed using the locus by locus phylogenetic approach (Table 2), so we used this more conservative approach to decide which strains to exclude from the final phylogenetic analysis.

**Table 2:**
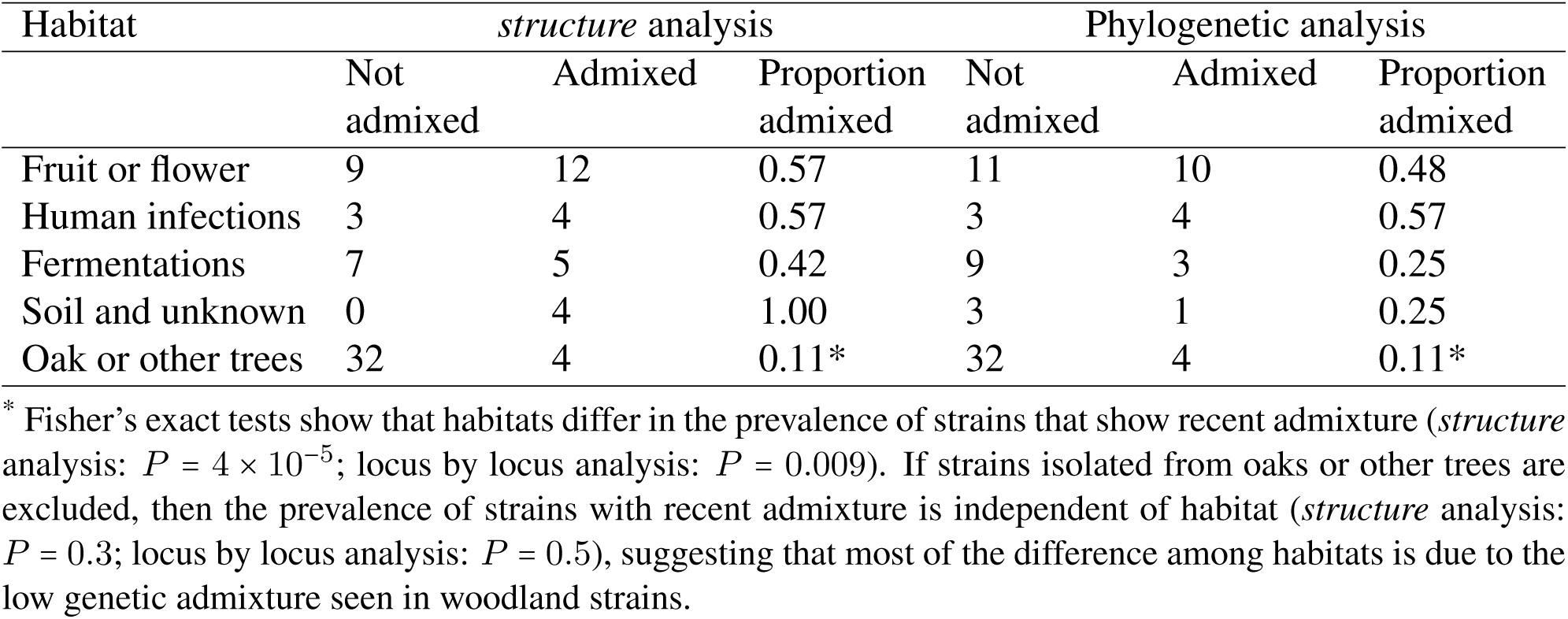
*S. cerevisiae* from trees in oak woodlands show less genetic admixture than those from other habitats.

| Habitat | *structure* analysis |  |  | Phylogenetic analysis |  |  |
| --- | --- | --- | --- | --- | --- | --- |
|  | Not admixed | Admixed | Proportion admixed | Not admixed | Admixed | Proportion admixed |
| Fruit or flower | 9 | 12 | 0.57 | 11 | 10 | 0.48 |
| Human infections | 3 | 4 | 0.57 | 3 | 4 | 0.57 |
| Fermentations | 7 | 5 | 0.42 | 9 | 3 | 0.25 |
| Soil and unknown | 0 | 4 | 1.00 | 3 | 1 | 0.25 |
| Oak or other trees | 32 | 4 | 0.11* | 32 | 4 | 0.11* |

\* Fisher's exact tests show that habitats differ in the prevalence of strains that show recent admixture (*structure* analysis: $P = 4 \times 10^{-5}$ ; locus by locus analysis: $P = 0.009$ ). If strains isolated from oaks or other trees are excluded, then the prevalence of strains with recent admixture is independent of habitat (*structure* analysis: $P = 0.3$ ; locus by locus analysis: $P = 0.5$ ), suggesting that most of the difference among habitats is due to the low genetic admixture seen in woodland strains.

## Results

### Seven genetically distinct populations of S. *cerevisiae*

We generated complete sequence data for whole centromeres from all 16 chromosomes of 47 *S. cerevisiae* strains from oak woodlands and fruit in the USA and Europe. We compared these sequences with a similar dataset previously described in Bensasson (2011) that includes 33 strains collected worldwide (Table 1). In *S. cerevisiae*, centromeres are small (up to 125 bp long), rapidly and neutrally evolving and have low recombination rates (Bensasson, 2011). By analyzing centromeres and their flanking DNA, every yeast chromosome is represented in our analysis. This strategy also minimizes the challenges to phylogenetic and population structure analysis presented by recombination and positive selection (Avise, 1994; Posada and Crandall, 2002; Brandvain *et al.*, 2014) that we would expect in a genome-wide analysis.

Using the Markov Chain Monte Carlo approach implemented in the program *structure* (Pritchard *et al.*, 2000; Falush *et al.*, 2003), we analysed data from 895 segregating sites in 13 kb of centromere sequence and identified five main populations in our sample of 80 strains (Figure 1a). These populations recapitulate those previously described in the world-wide sample of 33 strains: Malaysia, Sake, West Africa, Europe and the USA (Liti *et al.*, 2009). Although these five populations are also evident in our phylogenetic analysis of the same dataset there is not good statistical support for the European and USA clades. There do however appear to be well-supported distinct subpopulations of *S. cerevisiae* within both Europe and the USA (bootstrap values > 95%, Figure 2). These subpopulations are not identified by *structure* in any of the models we obtained, even when invoking a larger number of populations. We therefore used a hierarchical approach to test for population substructure within a subset of European strains (*N* = 23, Figure 1b) and within strains from the USA (*N* = 24, Figure 1c). This analysis revealed two subpopulations within Europe (Wine and European Oak), and two subpopulations within the USA (Pennsylvania Oak and North Carolina Oak). Overall, using this hierarchical *structure* approach, we identified a total of seven populations using *structure*, and these populations were also represented with well-supported clades in our phylogenetic analysis (Figure 2).

### A conservative test for recent genetic admixture

Using the seven populations identified above, we used two approaches to test for genetic admixture in each strain in our dataset. The first approach uses the industry standard software, *structure*, to estimate levels of genetic admixture given a panel of reference strains. We also developed a second locus by locus phylogenetic approach, which only invokes admixture for a strain if it has haplotypes at loci that are confidently assigned to differing known populations.

Analysis of admixture results for individual strains showed that 69 out of 80 strains were defined concordantly by the two methods (Supplemental File 1). In most of the remaining 11 cases, only *structure* invoked admixture in European strains (9 strains, Supplemental Files 1 and 2). Investigation of all 9 strains where only *structure* invokes admixture suggests that these are likely to be false positives because there was no strong or consistent phylogenetic support for their similarity to multiple populations (Supplemental Files 2 and 2). For some of these strains, *structure* appeared to invoke admixture to explain genetic divergence from known populations. For example, DBVPG1853m, a strain from North Africa is most similar to European populations of *S. cerevisiae*, yet it is also somewhat diverged from them (Figures 2 and 3). Locus by locus analysis showed that it is diverged from European strains, and most similar to the Wine or European populations (Supplemental Files 2 and 3). For this strain, *structure* invokes admixture between the Wine, European Oak and Sake populations however we see no evidence for greater similarity to European Oak or Sake than other populations at any single locus (Supplemental File 3).

In other cases, similarity between subpopulations can also lead to the potentially incorrect inference of admixture using the *structure* approach. For example, the wine strain RM11 consistently clustered with Wine or European strains at all loci where there was sufficient data to assign a population using the phylogenetic approach (12 loci, Supplemental Files 2 and 3). However, *structure* invoked admixture between the European Oak and Wine populations, even though there is no evidence that RM11 is more similar to the European Oak population than the Wine population at any loci (Supplemental File 3).

Two strains (SDO8s1 and SDO9s1) were defined as admixed using the locus by locus phylogenetic approach, but not using the *structure* approach. For both strains, we found strong evidence for admixture from the Wine population to the North Carolina Oak population at a single locus (CEN4, bootstrap support of at least 98%, Supplemental File 2). Using the *structure* approach we also detected some admixture from Wine to these two North Carolina Oak strains (4.7% and 5.5%), but these levels were below our threshold for defining admixture (1 locus out of 16, 6.25%). This analysis of the differences in admixture calls between the two methods suggests that the phylogenetic method is less likely to call false positives, while being more sensitive for the detection of low levels of admixture within a genome. We therefore used the admixture definitions from the locus by locus phylogenetic method for subsequent analyses.

### Genetic admixture is high in transient and low in stable habitats of *S. cerevisiae*

Because *S. cerevisiae* occurs in a broad range of habitats, we can use it as a model organism to test whether there is an effect of habitat on levels of genetic admixture. Fruit, flowers and insects represent habitats that are transient for yeast (Goddard and Greig, 2015) and *S. cerevisiae* also occurs in humans only transiently (Enache-Angoulvant and Hennequin, 2005). In contrast, trees are probably undisturbed for decades or even centuries. We therefore classified habitats in the worldwide sample of 80 strains according to whether they represent transient natural habitats such as (i) fruit or (ii) human infections, whether their provenance was less well defined from (iii) fermentations, or (iv) soil and unknown sources, or (v) whether they were sampled from well-defined stable woodland habitats (Table 2).

We found that the proportion of admixed strains is lower for strains from oak woodlands (11% of 36 strains) than for those of other habitats (Table 2) using both the *structure* (Fisher’s exact test, *P* = 4 × 10^−5^) and the locus by locus phylogenetic approaches (Fisher’s exact test, P = 0.009). When we excluded the *S. cerevisiae* from oaks or other trees, the proportion of admixed strains is similar across all habitats (59% of 44 strains, *structure* analysis: P = 0.3; 41% of 44 strains, locus by locus analysis: P = 0.5), suggesting that it is mainly in the woodland habitat that genetic admixture is peculiarly low (Table 2). Therefore our data suggest that levels of genetic admixture are high in transient and low in stable habitats (Table 2).

Many of the admixed *S. cerevisiae* strains defined using the conservative locus by locus phylogenetic method (8 out of 22 strains) showed a complex pattern of admixture involving at least three of the seven defined populations (Supplemental File 1). This suggests that the admixture we see in *S. cerevisiae* is not simply a consequence of recent hybridization resulting in the asexual descendants of F1 individuals. Furthermore, we were able to detect admixture only at a single locus for 8 strains, which suggests that in some cases backcrossing has occurred between hybrids and strains from single populations. All four admixed strains from woodlands showed admixture at a single locus, whereas admixture at only single loci occurred less often in strains from other habitats (4 out of 18 strains; Fisher’s exact test, *P* = 0.01). When strains showed admixture at only one locus, the admixture occurred at different loci for non-woodland strains but for the 4 strains from oak woodlands, they all showed evidence of admixture at the same locus (CEN4). Indeed, the CEN4 sequence was identical for 3 of these strains, suggesting that some of this admixture seen in oak strains does not result from independent events, and therefore that we could be overestimating the frequency of admixture in the oak habitat. Overall, the pattern of admixture observed suggests that the degree of admixture, as well as the frequency of admixture, could be lower in oak woodland habitats than in strains from other habitats.

### Distinct populations and low migration in woodland habitats

The inclusion of DNA sequences that show recent genetic admixture can lead to incorrect phylogenetic estimation, especially when admixture occurred recently between diverged populations (Posada and Crandall, 2002), as it does in this study. Thus, to better understand the true relationships between *S. cerevisiae* populations, it is necessary to reduce the effects of recent genetic admixture as much as possible. We therefore performed a phylogenetic analysis of the 58 strains in our dataset that did not show recent genetic admixture in our locus by locus phylogenetic analysis (Figure 3). Maximum likelihood phylogenetic reconstruction then showed that all but one of these 58 strains can be assigned to the seven known populations. This tree shows none of the long branch mosaic lineages reported in previous genome-wide analysis (Liti *et al.*, 2009), suggesting that our admixture filtering was successful.

One strain isolated from white teff grain in North Africa (DBVPG1853m) is not similar to any of the seven study populations. The bootstrap support in the analysis of pooled data from all loci (Figure 3) and in locus by locus analyses (Supplemental File 3) suggests that this strain may be a strain from an undersampled population that represents the closest outgroup to the Wine and European Oak populations of *S. cerevisiae*.

In addition, by overlaying habitat on our tree, phylogenetic analysis in the absence of admixture permits the identification of strains that have potentially migrated between habitats (Figure 3). Yeast strains from different oak woodlands mostly form distinct populations and differ from the strains of other habitats (Figure 3) and we found no evidence for the migration of yeast between oak woodland populations (Figure 3). However, there is evidence for the migration of yeast (i) from the North Carolina Oak population into a local vineyard, (ii) from the Wine population to oak trees, and (iii) from the Wine population to a medley of regions and substrates (Figure 3). Together, these observations suggest that strains from the Wine population migrate more between continents and habitats than the strains with woodland genotypes.

## Discussion

The wine yeast, *S. cerevisiae* has tremendous potential as a model for molecular ecology because it occurs naturally in several distinct habitats including fruit, flowers, insects and the bark of oak trees (Sniegowski *et al.*, 2002; Wang *et al.*, 2012; Hyma and Fay, 2013; Goddard *et al.*, 2010; Cromie *et al.*, 2013; Dashko *et al.*, 2016). *S. cerevisiae* is one of several model organisms whose genomes show evidence of recent genetic admixture from diverged populations (Liti *et al.*, 2009; Almeida *et al.*, 2015; Ludlow *et al.*, 2016; Hufford *et al.*, 2013; Brandvain *et al.*, 2014; Pool *et al.*, 2012; Sankararaman *et al.*, 2014). Therefore *S. cerevisiae* provides an excellent model to test an important question in molecular ecology: whether genetic admixture differs among habitats.

Using two approaches for defining admixed strains in a systematic and quantitative way, we show that patterns of genetic admixture differ between habitats (Table 2). Yeast strains from oak woodlands are less likely to show recent genetic admixture than those from other habitats (Table 2), and when it does occur the degree of admixture is lower and from fewer populations (Supplemental File 2 and 3). Consistent with our results, admixture has been noticed in the past for human-associated *S. cerevisiae* strains (Cromie *et al.*, 2013; Wang *et al.*, 2012) and for those fermenting cacao and coffee (Ludlow *et al.*, 2016), but there has been no compelling evidence reported previously for intraspecific admixture in yeast from oak woodlands.

By analyzing neutrally evolving centromeres with low recombination (Bensasson, 2011), we minimized the complications of recombination and natural selection on our analysis. Using only these centromere sequences, our results recapitulate genome-wide analyses that identified a second population of *S. cerevisiae* in Europe that represents the closest wild relatives to the *S. cerevisiae* Wine population (Cromie *et al.*, 2013; Almeida *et al.*, 2015), and the identification of two North American populations by Cromie *et al.* (2013). We were also better able to resolve the North American lineages using only centromere sequences compared to a previous analysis using genome-wide data (Liti *et al.*, 2009). Therefore our analysis of centromere sequences appears to have sufficient power to detect the lineages identified by genomic studies.

We developed a conservative locus by locus test to complement the use of *structure*, which is the standard Bayesian method used to estimate admixture (Pritchard *et al.*, 2000). A comparison of admixture calls using the two different approaches suggests that *structure* will sometimes invoke admixture to explain the divergence of a strain from defined populations, or it could incorrectly invoke admixture between genetically similar populations (Supplemental Files 1 and 3). Indeed, by using a locus by locus phylogenetic approach, we detect evidence for a distinct North African population (represented by DBVPG1853m in Figure 3) that was previously treated as an admixed strain in population genomic analyses (Liti *et al.*, 2009; Almeida *et al.*, 2015; Cromie *et al.*, 2013; Barbosa *et al.*, 2016). Molecular ecology and population genomic analyses of *S. cerevisiae* have mostly used only *structure* on data pooled across multiple loci to define admixed strains in order to better understand population structure in this species (Wang *et al.*, 2012; Almeida *et al.*, 2015; Barbosa *et al.*, 2016). Our analysis suggests the need for a more thorough investigation of admixture in yeast population genomic data using alternative methods.

When we removed admixed strains from our phylogenetic analyses, it became clear that woodland populations are distinct from one another, even when they occurred relatively close together in the Eastern USA (Figure 3). Given that we were unable to detect any migration (Figure 3) or admixture (Supplemental File 2) between woodland yeast populations it seems that yeasts in this ancestral habitat tend to be genetically isolated. Previous reports have suggested distinct oak-associated strains in the primeval forests of China (Wang *et al.*, 2012) and Brazil (Barbosa *et al.*, 2016). Our findings suggest that genetic isolation is not only a characteristic of Chinese and Brazilian forest populations, but that even strains from trees in Pennsylvania may be genetically isolated from trees in North Carolina.

Although we do not detect gene flow between oak woodlands, we do find evidence for both migration and admixture between the human-associated Wine population and woodland populations (Table 2, Supplemental Files 2 and 3). This supports past reports of migration of *S. cerevisiae* between vineyards and local oak trees (Goddard *et al.*, 2010; Hyma and Fay, 2013). It also mirrors the situation for *D. melanogaster* where human-associated populations show higher admixture than populations from non-urban regions, and there has been recent genetic admixture from cosmopolitan to ancestral populations (Pool *et al.*, 2012).

Transient habitats like fruit only exist for a few weeks and therefore must have been colonized recently by yeast (Goddard and Greig, 2015). Fruit flies, wasps, bees and other insects carry live *S. cerevisiae* in their guts and are therefore likely dispersal vectors for the migration of yeast to fruit (Goddard *et al.*, 2010; Stefanini *et al.*, 2012; Cromie *et al.*, 2013; Buser *et al.*, 2014). These insects visit multiple fruits and flowers, can fly long distances (Coyne and Milstead, 1987; Beekman and Ratnieks, 2000) and Drosophilids and honey bees at least have recently expanded to cosmopolitan distributions (Nunney, 1996; Whitfield *et al.*, 2006; Pool *et al.*, 2012). Insects associated with fruit can therefore potentially bring together *S. cerevisiae* strains from diverged populations. Furthermore, the spores of multiple strains of *S. cerevisiae* can survive passage through the guts of *Drosophila melanogaster* and the survivors are much more likely to undergo mating, and therefore admixture, than uneaten yeasts (Reuter *et al.*, 2007). Thus if yeast are primarily dispersed by insect vectors, yeasts from transient habitats are more likely to have cosmopolitan distributions and to show recent genetic admixture, as we observe here.

In contrast, oak tree bark is less nutrient rich (Goddard and Greig, 2015) and is therefore likely to attract fewer flying insects than rotting fruit. Consistent with this expectation, young oak trees have fewer yeast on their bark than older trees (Robinson *et al.*, 2016), suggesting that stable colonization of oak could be occurring over a period of years rather than weeks. Our observation of genetic isolation and low genetic admixture in oak woodland populations is therefore consistent with the lower migration distances and slower colonization expected for oak trees compared to fruit. It is also consistent with the lack of genetic admixture and the isolation by distance seen in *Saccharomyces paradoxus*, which is the closest relative of *S. cerevisiae* and has been studied almost exclusively from oak woodlands (Liti *et al.*, 2009). In addition, the degree of divergence that we observe between oak woodland populations may increase with geographic distance even in *S. cerevisiae:* North Carolina oak strains differ from Pennsylvania oak strains, while differing more from European oak strains (Figure 3)

*S. cerevisiae* is especially attractive as a model organism for molecular ecology and population genomics because of the resources already available for understanding its molecular evolution (Kellis *et al.*, 2003; Scannell *et al.*, 2011), molecular biology (Cherry *et al.*, 2011), experimental evolution (Rosenzweig and Sherlock, 2014), and for testing predictions in the laboratory (Cubillos *et al.*, 2009). Our study shows that the application of better methods for detecting genetic admixture in genomic data from woodland *S. cerevisiae* (Almeida *et al.*, 2015; Barbosa *et al.*, 2016) could lead to the generation of population genomic data that are unlikely to break the assumptions of most population genetic analyses. Therefore with more thorough testing for admixture and filtering, *S. cerevisiae* is likely to be an excellent model for population genomic analysis, despite its complex historical association with humans. There is some evidence that introgressions between species could confer adaptive traits in *S. cerevisiae* (Doniger *et al.*, 2008) and *Saccharomyces uvarum* (Almeida *et al.*, 2014). Population genomic analysis of intraspecific genetic admixture in maize revealed that gene flow from ancestral populations led to the adaptation of domesticated crops to the Mexican highlands (Hufford *et al.*, 2013). Given that *S. cerevisiae* is employed in a broad range of industries, including the production of wine, sake, beer, chocolate, and cacao, it will be especially interesting to apply new tools to study genome-wide patterns of admixture (Corbett-Detig and Nielsen, 2016) to reveal whether the genetic admixture seen among populations in *S. cerevisiae* plays a similar adaptive role in domestication.

## Acknowledgements

We thank Casey Bergman, Kelly Dyer, David Hall for helpful comments on the manuscript, and Stephanie Diezmann, Paul Sniegowski, Greg Wray, Anne Rouse, José Paulo Sampaio and Eladio Barrio for providing strains. This work was supported by the Natural Environ-ment Research Council through a NERC Fellowship to DB [grant number NE/D008824/1] (http://www.nerc.ac.uk).

## Authors’ Contributions

D.B. conceived and designed the research; V.T. performed the laboratory work; D.B. analysed the data and wrote the manuscript.

## Data Accessibility

DNA sequences determined for this study are available in GenBank: KT206234-KT206982. Perl scripts are available at https://github.com/bensassonlab/scripts, and locus by locus tree file data are available at https://github.com/bensassonlab/data. Most of the yeast strains used in this study are available from the National Collection of Yeast Cultures in the U.K. or the Portuguese Yeast Culture Collection.

## Supplemental Files

1. Supplemental File 1 in .pdf format showing the individual origin of the *S. cerevisiae* strains used and their genotypes.
2. Supplemental File 2 in tab-separated (.tsv) format that summarises the chromosome-by-chromosome analysis for each *S. cerevisiae* strain. Only clades with associated bootstrap values of at least 70% (7000 bootstrap replicates) were considered. Habitat descriptions are simplified versions of the categories shown in Table 2: “fermentation” includes wine, sake and other fermentations; “fruit” includes various fruit, flowers and insects; “human” means yeast were isolated from sites of human yeast infection; and “soilUnknown” describes habitat sources where a specific soil source is not well described or where the habitat source is completely unknown. Population assignments are given separately for each strain at each chromosome from 1 to 16. The column labeled “chrbychr70” shows the final genotype call for each strain in this study, and the column labeled “chrbychr70details” gives a summary of the different genotypes called for each strain.
3. Supplemental File 3 locus by locus trees in .pdf format showing the phylogenetic trees obtained for all 16 centromeres. Centromere loci are labelled CEN1 to CEN16, strains used as reference strains for their population are labelled with a coloured dot and strain names are coloured according to the final population assignment based on the locus by locus analysis of these trees: admixed strains are shown in black, blue is Malaysia, purple is European Wine, orange is West African, red is Sake, dark green with “YPS” prefix is Pennsylvania oak, mid-green with “SD”, “SM” or “ARN” prefix is North Carolina Oak, light green with a “ZP” prefix is European Oak. Trees were constructed from F84 distances using the neighbour joining method, and bootstrap values are percentages based on 10,000 bootstrap replicates. The phylogenies for every locus show that DBVPG1853m is most similar to European strains (10 out of 16 centromeres). At three loci, CEN2, 3 and, 5, there is also bootstrap support (at least 80% in all cases) that this strain represents the closest outgroup to the Wine and European Oak populations (Supplemental File 3).

